# Not all alloantibodies are created equal: IgG glycosylation and severity of antibody-mediated rejection in kidney transplantation

**DOI:** 10.1101/2025.05.28.656600

**Authors:** Johan Noble, Leandre M. Glendenning, Celine Dard, Anne Bourdin, Grace C. Carlson, Brian A. Cobb, Paolo Cravedi

**Affiliations:** Precision Immunology Institute, Translational Transplant Research Center (TTRC), Icahn School of Medicine at Mount Sinai, New York, NY, USA; Nephrology, Hemodialysis, Apheresis and Kidney Transplantation department, University hospital Grenoble, Grenoble, France; Univ. Grenoble Alpes, Inserm U 1209, CNRS UMR 5309, Team Cell Dynamics, Immunity, Metabolism & Cancer, Institute for Advanced Biosciences, Grenoble, France; Department of Pathology, Case Western Reserve University School of Medicine, Cleveland, OH, USA; Etablissement Français du sang, Grenoble-Alpes, 38000, France

**Author notes:** Co-first (Johan Noble, Leandre M. Glendenning). Co-senior (Brian A. Cobb, Paolo Cravedi). Corresponding authors: Brian Cobb, PhD, Case Western Reserve University School of Medicine 10900 Euclid Ave, Cleveland, OH 44106: Paolo Cravedi, MD, PhD, Icahn School of Medicine at Mount Sinai 1 Levy Place, New York, NY 10029.

**Keywords:** Antibody-Mediated Rejection, Glycosylation, Immunoglobulin G, Renal transplant

## Abstract

**Introduction:** Antibody-mediated rejection (AMR) is a leading cause of kidney transplant (KT) failure, driven by donor-specific anti-HLA antibodies (DSA). However, not all patients with DSA experience accelerated graft loss, suggesting that factors beyond antibody presence influence AMR severity. Post-translational modifications, particularly glycosylation of Immunoglobulin-G (IgG), play a critical role in modulating antibody function. This study investigates the association between IgG glycosylation profiles and the risk and severity of AMR in KT recipients.

**Methods:** We prospectively analyzed 65 KT patients, including 26 with acute AMR (aAMR), 27 with chronic-active AMR (caAMR), and 12 controls without rejection. IgG glycosylation was quantified using lectin-based ELISA, focusing on mannose, fucose, sialic acid, and bisecting N-acetylglucosamine (GlcNAc) levels.

**Results:** Results showed that bisecting GlcNAc levels of total IgG were significantly higher in caAMR patients than controls (p=0.019) and aAMR patients (p=0.045). Multivariable analysis revealed that higher bisecting GlcNAc levels of IgG were independently associated with glomerulitis [g-score, OR: 2.7 (95%CI: 1.2-6.7), p=0.019] and chronic glomerulopathy [cg-score, OR: 2.8 (95%CI: 1.3-7.5), p=0.021], independent of DSA presence.

**Conclusions:** These findings indicate an association between IgG glycosylation, particularly bisecting GlcNAc, and AMR severity. IgG glycosylation profiles could serve as biomarkers for AMR risk and severity, offering new insights into the mechanisms of AMR and potential therapeutic targets.

## Introduction

Antibody-mediated rejection (AMR) is a major cause of kidney transplant loss.^1^ Although the pathophysiology of AMR is not fully understood, donor-specific anti-HLA antibodies (DSA) are thought to play an important pathogenic role. When present, these antibodies target endothelial graft cells, leading to their direct activation,^2^ the initiation of the complement cascade, and the recruitment of immune cells.^3,4^ Antibodies can also promote endothelial injury through antibody-dependent cellular cytotoxicity (ADCC) and the release of pro-inflammatory cytokines, both events that are primarily mediated by natural killer (NK) cells.^5,6^

While DSA are central to AMR pathogenesis - triggering complement-dependent cytotoxicity (CDC), antibody-dependent cellular cytotoxicity (ADCC), and proinflammatory injury - not all patients with DSA experience graft loss. This disconnect suggests that antibody quantity alone fails to explain clinical outcomes.

Evidence has shown that post-translational modification of antibodies affects their function.^7,8^ The type and level of antibody glycosylation can influence its ability to activate complement and induce ADCC, as well as their stability and circulating half-life.^9^ Glycosylation of Immunoglobulin G (IgG) involves the addition of carbohydrate at a conserved site in the Fc region (Asn297), and, less frequently, in the Fab region.^10^ Glycan sites are primarily introduced during somatic hypermutation. The N-linked glycosylation starts with the transfer of a high mannose glycan from a dolichol-glycan donor in the endoplasmic reticulum (ER) and is critical for protein folding quality control (**Supplementary Figure 1A**). Upon exit from the ER and entry into the Golgi apparatus, the high mannose structures are matured by sequential removal of all but three mannoses, leaving a pentasaccharide core (**Supplementary Figure 1B**), and then rebuilt through the addition of N-acetylglucosamine (GlcNAc) residues (**Supplementary Figure 1C**). This results in complex N-glycans carrying varied amounts of GlcNAc branch points, core fucosylation, galactose and sialic acid residues. Glycan structures found on IgG are typically limited to bi-antennary with and without a bisecting GlcNAc (**Supplementary Figure 1D**), a branching modification that influences IgG function and enhances Fc receptor binding. Functionally, sialylation (addition of terminal N-acetylneuraminic acid with an α2,6 linkage to the underlying galactose) and core fucosylation (fucose attached with an α1,6 linkage to the asparagine-proximal GlcNAc) are associated with reduced ADCC properties by reducing Fc gamma receptors (FcγR)III affinity.^11^

IgG glycosylation profile has been associated with changes in the severity of various autoimmune diseases.^12^ A study in kidney transplant recipients showed that low sialylation and high bisecting GlcNAc of anti-donor specific HLA antibodies (DSA) was associated with higher risk of AMR.^13^ However, this study assessed only post-translational modifications in IgG3, Herein, we hypothesized that total IgG glycosylation profile may be associated with the risk and the severity of both AMR and caAMR.

## Materials

### Study population

We prospectively included kidney transplant recipients from a single center (Grenoble-Alpes University, France) between January 2016 and May 2024 with a biopsy-proven AMR and control kidney transplant patients with a per-protocol biopsy showing no signs of microvascular inflammation (MVI). All biopsies were reviewed by a single pathologist who classified the biopsies according to Banff 2022.^14^ In brief, AMR was defined by the presence of MVI, i.e. glomerulitis (g) and peritubular capillaritis (ptc). If the MVI score was 2 or above, AMR was confirmed by the presence of C4d deposition in peritubular capillaries and vasa recta and/or by the presence of DSA. If other features of AMR, such as acute thrombotic microangiopathy were present, but the MVI score was below 2, AMR was confirmed by the presence of C4d deposition, regardless of DSA status. C4d deposition was quantified using anti-C4d antibody on formalin-fixed, paraffin-embedded sections. Chronic active AMR (caAMR) was defined by the presence of acute lesions (including C4d) and chronic lesions (transplant glomerulopathy (cg) and or peritubular capillary multilayering).

All patients signed an informed consent form. All medical data were collected from our database [CNIL (French National committee for data protection) approval number 1987785v0].

### Anti-HLA measurement

Anti-HLA antibodies were measured at the time of biopsy in all patients using One Lambda-Thermofisher Labscreen, Class I and Class II. Antibody specificity was identified using a Luminex® 200 Instrument (Luminex Corporation, Austin, TX). The MFI threshold of positivity for an anti-HLA and /or a DSA was 500. The sera used for the post-translational analysis were leftovers from the anti-HLA analysis at the Etablissement Français du Sang (EFS) of Grenoble, HLA department.

### Immunoglobulin G purification

We purified total serum IgG using Protein A IgG Purification Kit (Pierce^TM^, Thermo Scientific^TM^, CAT: 44667) and following manufacturer’s instructions. Briefly, the procedure consisted of passing the sera sample into the column and then eluting the IgG bound to the resin using an elution buffer (pH 2.8). For each sample, IgG concentration was measured by absorbance at 280 nm and purity was confirmed using SDS-PAGE.

### Quantification of glycosylation by Lectin ELISA

Purified IgG were diluted to 1ng/μL in PBS, plated into a 96-well high-binding ELISA plate (Microlon High Binding; Greiner BioOne) and incubated overnight at 4 °C. The plate was blocked with carbohydrate free blocking solution (Vector Labs) for 1 hour at room temperature. Biotinylated lectins (Vector Labs) were diluted to 1μg/mL in carbohydrate-free blocking solution and incubated for 1 hour at room temperature. Different biotinylated lectins were used: Sambucus nigra agglutinin (SNA): binds to α2,6-linked Neu5Ac and is used to quantify the sialylation of IgG. Concanavalin A (ConA): binds to high mannose and hybrid N-glycans and correlates with more immature IgG. Lens culinaris agglutinin (LCA): binds to core fucosylated N-glycans. Phaseolus vulgaris Erythroagglutinin (PHA-E) binds to bisecting GlcNAc to quantify bisecting GlcNAc on IgG. Anti-Fc measures the amount of immobilized IgG to normalize the results. The amount of IgG coated was 0, 100ng and 200ng to assess saturation of the signal and we thereafter used the results of 100ng for consistency. Signal was detected using europium-conjugated streptavidin (Perkin Elmer) and time-resolved fluorescence as measured in a Victor Nivo plate reader.

### Statistical analysis

Normally distributed quantitative variables were expressed as mean ± standard deviation (SD). Non-normally distributed quantitative variables were expressed as median [interquartile ranges (IQR)]. All glycosylation values were non-normally distributed (p-value from Shapiro-Wilk Test <0.005). Qualitative variables were expressed as numbers and percentages. For statistical comparison between multiple groups, we used Kruskal-Wallis test. For statistical comparisons between two groups, we used unpaired Mann-Whitney U Tests. The chi-squared test was used to compare categorical data.

To identify factors associated with each ordinal outcome variable (i.e., scores of Banff components), we employed ordinal logistic regressions using the proportional odds model using the *clm* function from the ordinal R package (R version 4.4.2, https://www.r-project.org). Variables with a p-value < 0.05 in univariate analyses were included in the multivariable ordinal logistic regression models. The presence of DSA was included in the multivariable model regardless of its univariate significance, based on their clinical relevance. To assess the validity of the proportional odds assumption, we conducted the Brant test. Statistical significance was defined as a two-sided p-value < 0.05.

## Results

### Baseline characteristics of the study population

The study included 65 kidney transplant recipients, 26 (40%) with biopsy-proven aAMR, 27 (41.5%) with caAMR, and 12 controls with no histological signs of rejection (**Table 1**). There was no difference between aAMR, caAMR and controls in the main characteristics such as the age at the time of biopsy, gender, previous transplantation number, level of sensitization, type of donor, HLA mismatches, history of diabetes, and induction therapy. The median time between the transplantation and the biopsy was 0.7 years [0.1 – 3.4] in the aAMR group, 4.6 [2.9 – 14.1] for the caAMR group, and 0.3 [0.3 – 0.3] for the control group, p <0.001. The immunosuppressive regimen consisted of tacrolimus, mycophenolate mofetil, and steroids for all patients.

**Table 1:** Baseline characteristics of study participants.

|  | **CTL**<br>**(N = 12)** | **aAMR**<br>**(N = 26)** | **caAMR**<br>**(N = 27)** | ***p-value*** |
| --- | --- | --- | --- | --- |
| Age - years | 49.1 ± 14 | 44.1 ± 15 | 41.4 ± 16 | 0.522 |
| Gender M/F ratio | 3 | 0.9 | 1.8 | 0.310 |
| History of previous transplantation – N (%) | 0 | 6 (23.0) | 3 (11.1) | 0.200 |
| Nephropathy – N (%) |  |  |  | 0.522 |
| Diabetes | 0 (0.0%) | 0 (0.0%) | 1 (3.7%) |  |
| Drug-related | 0 (0.0%) | 0 (0.0%) | 1 (3.7%) |  |
| FSGS | 0 (0.0%) | 3 (11.5%) | 1 (3.7%) |  |
| Genetic | 1 (8.3%) | 2 (7.7%) | 0 (0.0%) |  |
| IgAN | 1 (8.3%) | 3 (11.5%) | 1 (3.7%) |  |
| Unknown | 6 (50.0%) | 7 (26.9%) | 14 (51.9%) |  |
| Auto-immune disease | 0 (0.0%) | 3 (11.5%) | 0 (0.0%) |  |
| TMA | 0 (0.0%) | 1 (3.8%) | 1 (3.7%) |  |
| Hypertensive nephropathy | 2 (16.7%) | 3 (11.5%) | 0 (0.0%) |  |
| Acute interstitial nephropathy | 0 (0.0%) | 2 (7.7%) | 4 (14.8%) |  |
| Acute tubular necrosis | 0 (0.0%) | 0 (0.0%) | 1 (3.7%) |  |
| Polycystic disease | 2 (16.7%) | 2 (7.7%) | 3 (11.1%) |  |
| Delay of biopsy - months | 0.3 [0.3 – 0.3] | 0.7 [0.1 – 3.3] | 4.6 [2.9 – 14.1] | <0.001 |
| PRA (%) | 24.2 ± 39 | 54.6 ± 45 | 34.6 ± 46 | 0.223 |
| Living donor – N (%) | 1 (12.5) | 6 (24.0) | 5 (20.0) | 0.011 |
| Donor age - years | 49.8 ± 16 | 47.9 ± 16 | 44.1 ± 15 | 0.643 |
| HLA mismatch A/B/DQ - number | 6.5 [5 – 7] | 6.0 [5 – 7] | 5.5 [5 – 7] | 0.356 |
| History of diabetes – N (%) | 0 | 0 | 1 (4.8) | 0.471 |
| Anti-thymoglobulin induction – N (%) | 9 (75) | 24 (92.3) | 26 (96.3) | 0.754 |
| MVI ≥ 2 – N (%) | 0 | 23 (88.5) | 26 (96.3) | <0.001 |
| C4d positivity– N (%) | 0 | 19 (73.0) | 9 (33.3) | <0.001 |
| cg-score positivity– N (%) | 0 | 0 | 25 (96.2) | <0.001 |
| sCreat at the time of biopsy – mg/dL | 1.74<br>[1.3 – 2.3] | 1.59<br>[1.1 – 2.5] | 2.04<br>[1.5 – 3.1] | 0.628 |
| Proteinuria at rejection – g/g | 0.2 [0.1 – 0.6] | 0.3 [0.2 – 0.7] | 0.8 [0.3 – 1.8] | 0.928 |

FSGS: Focal segmental glomerulosclerosis; HLA: Human Leukocyte Antigen; IgAN: IgA nephropathy; MVI: microvascular inflammation; PRA: panel reactivity antibody; sCreat: serum creatinine; TMA: thrombotic microangiopathy.

Twenty-one aAMR patients (80.8%) had donor-specific antibody (DSA) at the time of the biopsy, compared to 20 (74.1%) in the caAMR group (p=0.560). None of the patients in the control group had DSA. The number of DSA was also similar between the two rejection groups (1.6±1.0 vs 1.4±1.0, p=0.727) and most of DSA were class II (81% versus 85%, p=0.731) for aAMR and caAMR, respectively. The mean fluorescence intensity (MFI) of the immunodominant DSA was also similar (8113 ± 6506 vs 5064 ± 6083, p=0.273). The median follow-up period post-transplantation was 2.8 [2.0 – 4.2] years in the aAMR group, 8.4 [4.0 – 11.2] years in the caAMR group, and 1.2 [0.4 – 2.4] years in the control group, p<0.001.

### Associations between IgG glycosylation and the risk of AMR

The quantification of IgG glycan composition (relative amounts of mannose, fucose, sialic acid, and bisecting GlcNAc) was assessed using the ConA, LCA, SNA, and PHA-E lectins, respectively, and normalized with the Fc levels (**Figure 1**). We found a positive correlation between mannose, core fucose, α2,6-linked sialic acids. In contrast, bisecting GlcNAc were not significantly associated with the other glycoforms (**Supplementary Figure 2)**.

**Figure 1:**
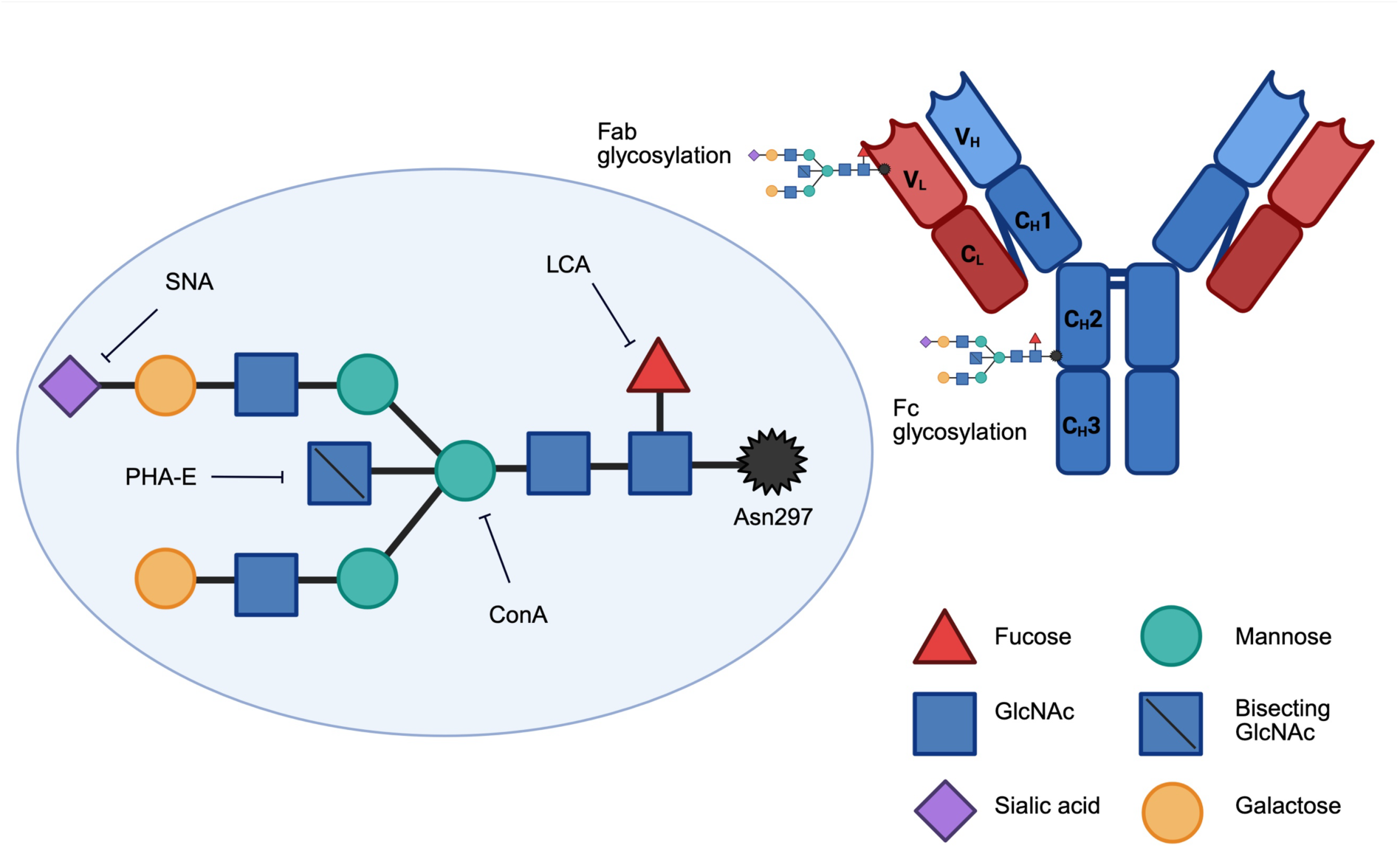
Representation of Immunoglobulin G glycosylation sites and detection. The antibody regions are labeled as C_H_1, C_H_2, C_H_3 (constant heavy domains), V_H_ (variable heavy), V_L_ (variable light), and C_L_ (constant light). A critical glycosylation site at *Asn297* in the Fc region is shown, along with Fab glycosylation. The figure also illustrates key glycan components, including fucose, sialic acid, mannose, bisecting GlcNAc, and galactose. Additionally, lectins with binding specificity (LCA, PHA-E, SNA, ConA) are indicated, suggesting their utility in glycan analysis.

We first compared the different IgG glycoforms among the 3 groups. The amounts of mannose (ConA/Fc), core fucose (LCA/Fc), and bisecting GlcNAc (PHA-E/Fc) were significantly lower in the control group than in the caAMR group. Bisecting GlcNAc (PHA-E/Fc) was significantly higher in the caAMR and aAMR groups compared to controls, and significantly higher in caAMR compared to aAMR patients (**Figure 2A**). Core fucose (LCA/Fc) was significantly higher in both caAMR and aAMR groups compared to controls (**Figure 2B**). Mannose (ConA/Fc) was significantly higher in caAMR than in controls (**Figure 2C**). There was no statistical difference among the 3 groups in the α2,6-sialylation (SNA/Fc) levels (**Figure 2D**).

**Figure 2:**
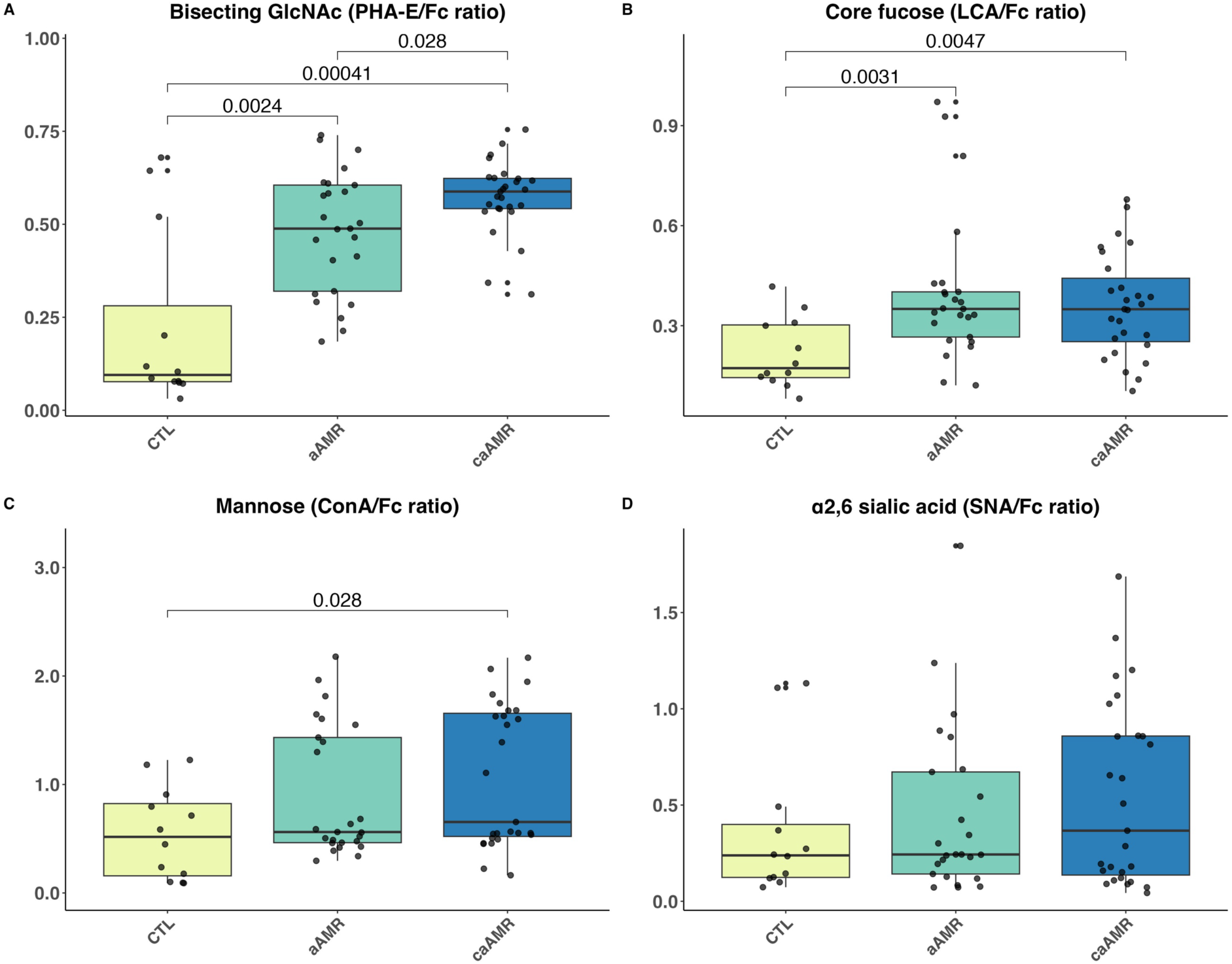
Associations between IgG post-translational modifications and the presence of antibody-mediated rejection. Box plots comparing the amounts of bisecting GlcNAc (PHA-E/Fc ratio in panel A), core fucose (LCA/Fc ratio in panel B), mannose (ConA/Fc ratio in panel C), and α2,6-sialylation (SNA/Fc in panel D) among control patients, patients with acute antibody-mediated rejection and chronic-active antibody mediated rejection. p-values < 0.05 is indicated for each comparison. aAMR: acute Antibody-Mediated Rejection; caAMR: chronic active Antibody-Mediated Rejection; CTL: control patients, ConA: Concanavalin A; LCA: Lens culinaris agglutinin; SNA: Sambucus nigra agglutinin; PHA-E: Phaseolus vulgaris Erythroagglutinin.

### Associations between IgG glycosylation and the severity of rejection

We then compared the amount of each glycoform with the severity of AMR using each of the AMR parameters listed in the Banff score (0 to 3), i.e. g-, cg-, and ptc-scores and C4d deposition intensity (0 to 3). We used cg-score to quantify the chronicity of lesions. We found no association between bisecting GlcNAc (PHA-E/Fc) and g-score severity (**Figure 3A**). We found that mannose (ConA/Fc), α2,6-sialic acids (SNA/Fc) and core fucose (LCA/Fc) staining was increased in samples from patients with the highest g-scores compared to those with lower scores (**Figure 3B-D**). Within AMR cases, core fucosylation was lower in patients with less severe glomerulitis (g-score = 0) than in those with more pronounced inflammation (**Figure 3B**). Regarding the other Banff scores, we found that a higher cg-score was associated with increased sialylation (SNA/Fc; **Supplementary Figure 3**).

**Figure 3:**
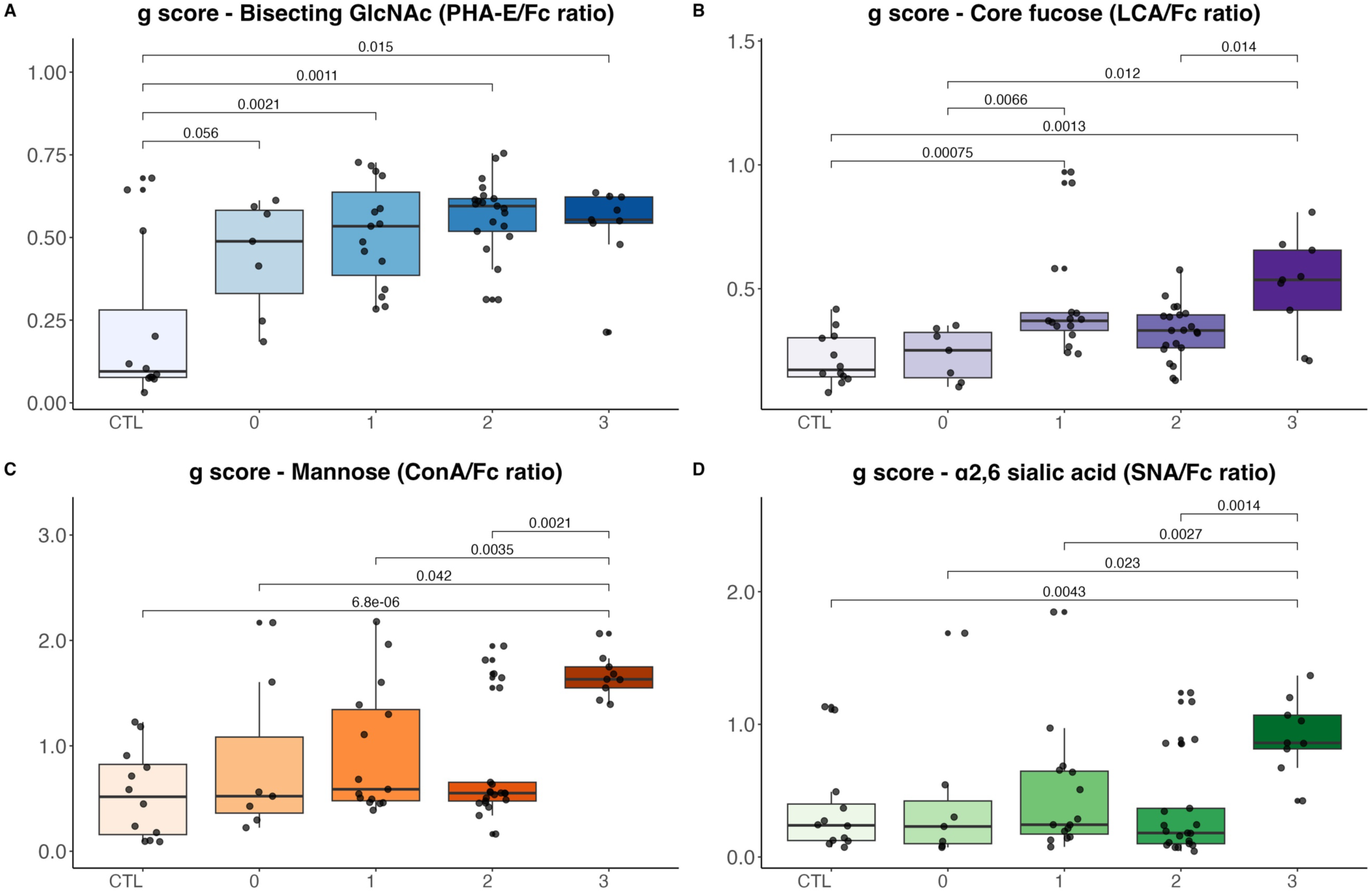
Associations between IgG post-translational modifications and the severity of rejection. Box plots comparing the levels of bisecting GlcNAc (PHA-E/Fc ratio in panel A), core fucose (LCA/Fc ratio in panel B), mannose (ConA/Fc ratio in panel C), and α2,6-sialylation (SNA/Fc in panel D) with the severity score of glomerulitis (g-score). p-values < 0.05 is indicated for each comparison. ConA: Concanavalin A; cg: chronic glomerulopathy; c4d: complement factor 4d; g: glomerulitis; LCA: Lens culinaris agglutinin; PHA-E: Phaseolus vulgaris Erythroagglutinin; ptc: peritubular capilaritis; SNA: Sambucus nigra agglutinin.

### Association between IgG glycosylation and DSA

Not all patients in our cohorts had DSA. Therefore, we tested the association between IgG glycosylation and DSA. Among the different glycosylation tested, we found that only bisecting GlcNAc was associated with the presence of DSA in patients with AMR. In particular, DSA positive patients had higher levels of bisecting GlcNAc (**Figure 4**).

**Figure 4:**
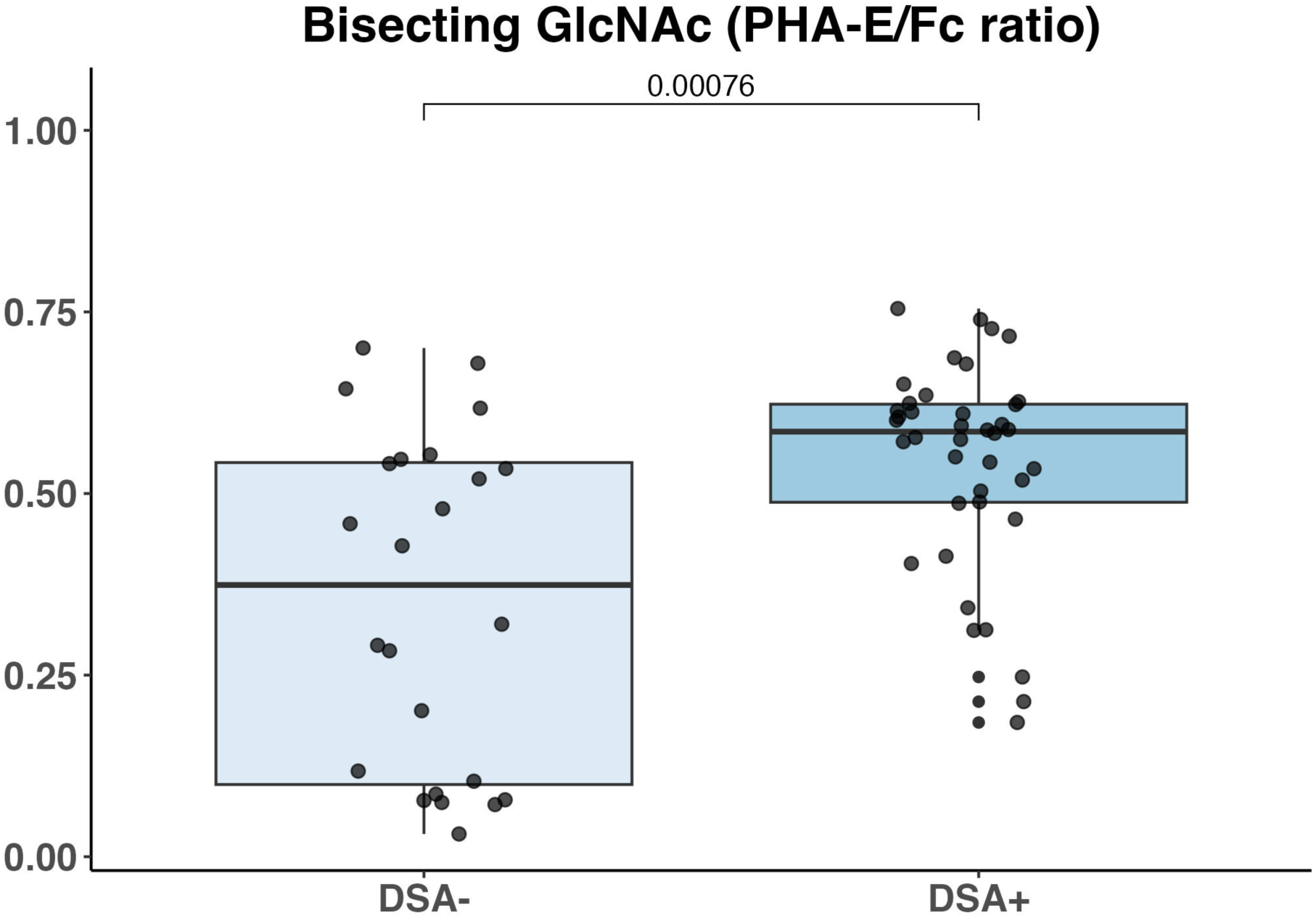
N-glycan bisecting GlcNAc (PHA-E/Fc) IgG glycans in patients with or without DSA at the time of biopsy. DSA: Donor-specific antibody; PHA-E: Phaseolus vulgaris Erythroagglutinin.

We repeated the analyses comparing glycoform levels in controls, aAMR, and caAMR cohorts, by stratifying patients based on the presence of DSA. In DSA negative patients, aAMR and caAMR remained significantly higher than controls for bisecting GlcNAc and core fucose (**Supplementary Figure 4A-B**). In DSA positive patients, bisecting GlcNAc was higher in caAMR compared to aAMR patients (**Supplementary Figure 4A**). Mannose and α2,6-sialylation were no longer significantly different when we stratified patients based on DSA status (**Supplementary Figure 4C-D**).

### Multivariable analysis of factors associated with histologic features of AMR

Finally, we tested whether the IgG glycosylation and AMR severity are independently associated. We started testing an association between all clinical variables, IgG glycosylation, presence of DSA, and histological features of AMR using a univariable ordinal regression model. The variables with a p-value < 0.05 in univariate analyses were considered for inclusion in the multivariable ordinal logistic regression model using AMR histological scores as ordinal independent variables (**Table 2**). We found that the g-score was significantly associated with the presence of DSA and higher bisecting GlcNAc (PHA-E/Fc) levels and Pts-score with the presence of DSA and core fucose. In contrast, cg-score was associated with bisecting GlcNAc, α2,6 sialic acid levels, and time after transplant, but not with the presence of DSA (**Table 2**).

**Table 2:** Multivariable ordinal regression analysis of the association between clinical variable, IgG post-translational modifications, and AMR severity scores in the overall cohort.

|  | g-score |  | ptc-score |  | cg-score |  |
| --- | --- | --- | --- | --- | --- | --- |
|  | OR<br>[95%CI] | p-value | OR<br>[95%CI] | p-value | OR<br>[95%CI] | p-value |
| Presence of DSA | **5.20**<br>**[1.2–25.1]** | **0.032** | **3.59**<br>**[1.2–11.6]** | **0.028** | 0.83<br>[0.2–2.9] | 0.768 |
| Bisecting GlcNAc | **2.72**<br>**[1.2–6.7]** | **0.019** | 1.38<br>[0.7–2.4] | 0.263 | **2.81**<br>**[1.3–7.5]** | **0.021** |
| Core fucose | 5.62<br>[0.8–43.9] | 0.088 | **2.24**<br>**[1.3–3.9]** | **0.003** |  |  |
| α2,6 sialic acid | 0.32<br>[0.05–1.7] | 0.184 | | | **2.04**<br>**[1.2–3.7]** | **0.014** |
| Mannose | 1.44<br>[0.7–3.2] | 0.354 |  |  |  |  |
| Delayed graft function | 0.16<br>[0.01–1.7] | 0.161 |  |  |  |  |
| Time between KT and biopsy (yr) |  |  |  |  | **1.11**<br>**[1.0–1.2]** | **0.012** |

Bisecting GlcNAc (PHA-E/Fc ratio); Core fucose (LCA/Fc ratio); α2,6 sialic acid (SNA/Fc ratio); mannose (ConA/Fc ratio).

## Discussion

In this study, we found an association between total IgG core-fucosylation, bisecting GlcNAc, and detectable mannose, and the presence of biopsy-proven AMR in kidney transplant recipients. IgG α2,6-sialylation was not associated with AMR in our cohort. We also found that bisecting GlcNAcylation was significantly higher in patients with caAMR versus aAMR in patients with a circulating DSA at the time of AMR diagnosis compared to DSA negative and C4d negative AMR. Moreover, bisecting GlcNAcylation was significantly associated with glomerulitis and chronic glomerulopathy lesions. Interestingly, the significant association between bisecting GlcNAcylation and g and cg-scores lesions was independent of the presence of DSA.

Our new findings confirm and expand results from prior smaller studies.^15^ Pernin *et al.* performed mass spectrometry on isolated IgG3 *de novo* DSA and showed that higher bisecting GlcNAc glycosylation was higher in AMR patients overall compared to patients DSA+, but without histological AMR.^13^ If mass spectrometry allows a precise characterization of glycoforms, it can also exhibit higher technical variability compared to other methods, potentially affecting reproducibility.^16^ ELISA, on the contrary, is well-suited for large-scale studies due to its simplicity and ability to process many samples simultaneously, is generally less expensive and requires less specialized equipment compared to mass spectrometry, and can be standardized across different laboratories, enhancing reproducibility.^17^

Our findings are in line with the literature about bisecting GlcNAc. Bisecting GlcNAcylation levels can modify the Fc activity of IgG by influencing its binding affinity to FcγRs. Specifically, the presence of a bisecting GlcNAc moiety has been shown to inhibit Fc core-fucosylation and enhance the binding of Fc to FcγRIIIa, an activating Fcγ receptor.^18,19^ This enhancement in binding affinity can lead to increased ADCC activity, which conceivably results in increased glomerular injury induced by HLA and non-HLA antibodies.

In the study by Pernin *et al.,* sialylation of IgG3 DSA was found to be associated with a reduced risk of AMR. Sialylation of total IgG assessed at the time of transplantation was also associated with reduced AMR risk during the follow-up in another study.^20^ In our cohort, we did not find any correlation between IgG sialylation and the presence of AMR. Together, these data suggest a reduced activity of HLA and non-HLA antibodies in kidney transplantation when associated with a higher sialylation. This hypothesis is supported by the literature, as sialylation of the IgG Fc domain has been shown to impair complement-dependent cytotoxicity (CDC) by reducing the binding of C1q,^21^ decreasing ADCC (in the presence of core fucosylation) by reducing the binding affinity of IgG to FcγRIIIa,^22^ and exhibiting anti-inflammatory effects.^23^

The significance of our data extends to individuals with autoimmune diseases, where alterations in IgG glycosylation have been well-documented. For instance, in rheumatoid arthritis, there is a notable decrease in galactosylation and sialylation, and increased fucosylation in IgG during active phases of the disease,^24^ suggesting that these glycosylation changes can serve as biomarkers for disease activity and therapeutic response. Similarly, systemic lupus erythematosus patients exhibit decreased IgG galactosylation and sialylation, increased bisecting GlcNAc, and decreased core fucose. ^25,26^ These glycosylation changes are associated with disease activity and specific clinical manifestations, such as nephritis and the onset of antinuclear antibodies. IgG glycosylation patterns, including sialylation, bisecting GlcNAcylation, fucosylation, and mannosylation, are significantly altered in various other autoimmune and inflammatory diseases, and these changes have been proposed as potential biomarkers for disease activity and severity.^27,28^

One limitation of our study is that we analyzed total IgG glycosylation, instead of DSA-specific IgG glycosylation, as they may theoretically differ. However, Sonneveld *et al.* demonstrated that the Fc-glycosylation profiles of total serum IgG1 and IgG3 were highly similar to those of antigen-specific anti-K antibodies.^29^ In that study, the authors analyzed the Fc-tail glycosylation of both total serum and antigen-specific IgG1 and IgG3 using mass spectrometry and found that the glycosylation patterns were consistent across both types, with little individual variability. Using an integrated method for simultaneous protein quantitation and glycosylation profiling of antigen-specific IgG1, Gijze *et al.* also found that the glycosylation profiles of antigen-specific IgG1 are comparable to those of total IgG1 in a large coronavirus vaccination cohort.^30^ Of note, we found that the differences in bisecting GlcNAcylation persist in patients with and in those without DSA, suggesting that, although DSA may have a unique glycosylation profile, some key features are probably shared across antibody specificities, but this will require further investigation.

In conclusion, we found a significant, independent correlation between total IgG glycosylation and AMR in kidney transplant recipients. More specifically, we showed the association of IgG bisecting GlcNAc regardless of the presence of DSA, with glomerulitis and chronic glomerulopathy in graft biopsy. Therefore, our findings provide the rationale for larger studies testing the hypothesis that IgG glycosylation can be used as a biomarker for the risk of AMR and caAMR. Strategies to change IgG glycosylation may also be leveraged therapeutically to improve the outcomes of transplant recipients.

## Disclosure

The authors of this manuscript have no conflicts of interest to disclose.

## Acknowledgments

We thank Shivani Wadnerkar and Friederike Selbach for helping isolate IgG from patients’ sera.

## Author contribution

JN isolated IgG from patients’ sera, performed the statistical analysis, and wrote the initial draft of the manuscript. LMG and GCC measured IgG glycosylation; CD and AB collected the sera samples and the patient’s data; PC and BAC had the original idea, supervised the work, and revised the manuscript.

## Data availability statement

Data are available on reasonable request from the corresponding authors.

## Supplementary Material

Supplementary Figures

## Abbreviations

AMR: Antibody-mediated rejection
ADCC: Antibody-dependent cellular cytotoxicity
ConA: Concanavalin A
DSA: Donor-specific Antibody
g: glomerulitis
GlcNAc: N-acetylglucosamine
IgG: Immunoglobulin G
LCA: Lens culinaris agglutinin
ptc: peritubular capilaritis
MVI: microvascular inflammation
Neu5Ac: sialic acid
SNA: Sambucus nigra agglutinin

