## supplemental material for "Not all alloantibodies are created equal: IgG glycosylation and severity of antibody-mediated rejection in kidney transplantation"

### SUPPLEMENTARY MATERIALS

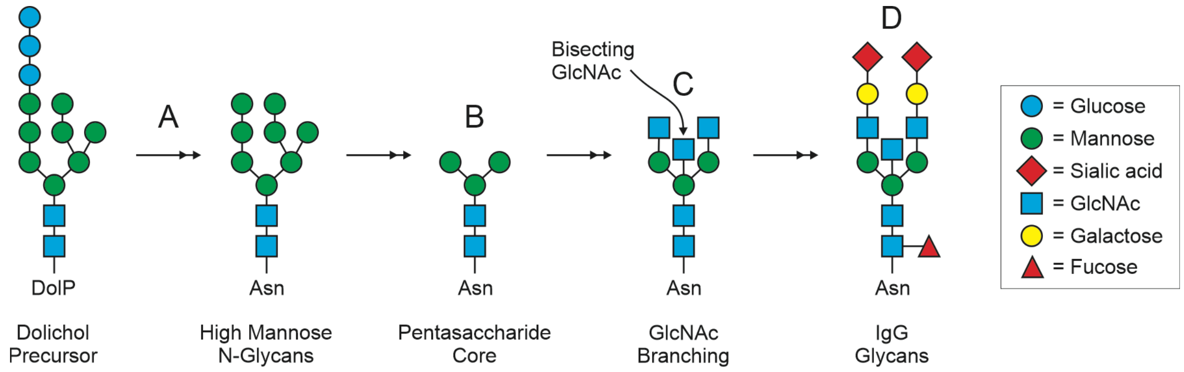


**Supplementary Figure 1: Key Structural Components and Modifications in N-Glycan Biosynthesis.** Illustration of the essential structural elements and modifications involved in the biosynthesis of N-glycans. Panel A highlights the dolichol phosphate (DolP) and its precursor role in the initial stages of glycan assembly and depicts high-mannose N-glycans attached to an asparagine (Asn) residue, representing an early intermediate in the pathway. Panel B shows the pentasaccharide core, a conserved structure in N-glycan synthesis. Panel C shows the branching process, marked by the addition of N-acetylglucosamine (GlcNAc) to the core. Panel D showcases the diversity of terminal sugar modifications, including glucose, mannose, sialic acid, GlcNAc, galactose, and fucose, which contribute to the functional variability of mature N-glycans.

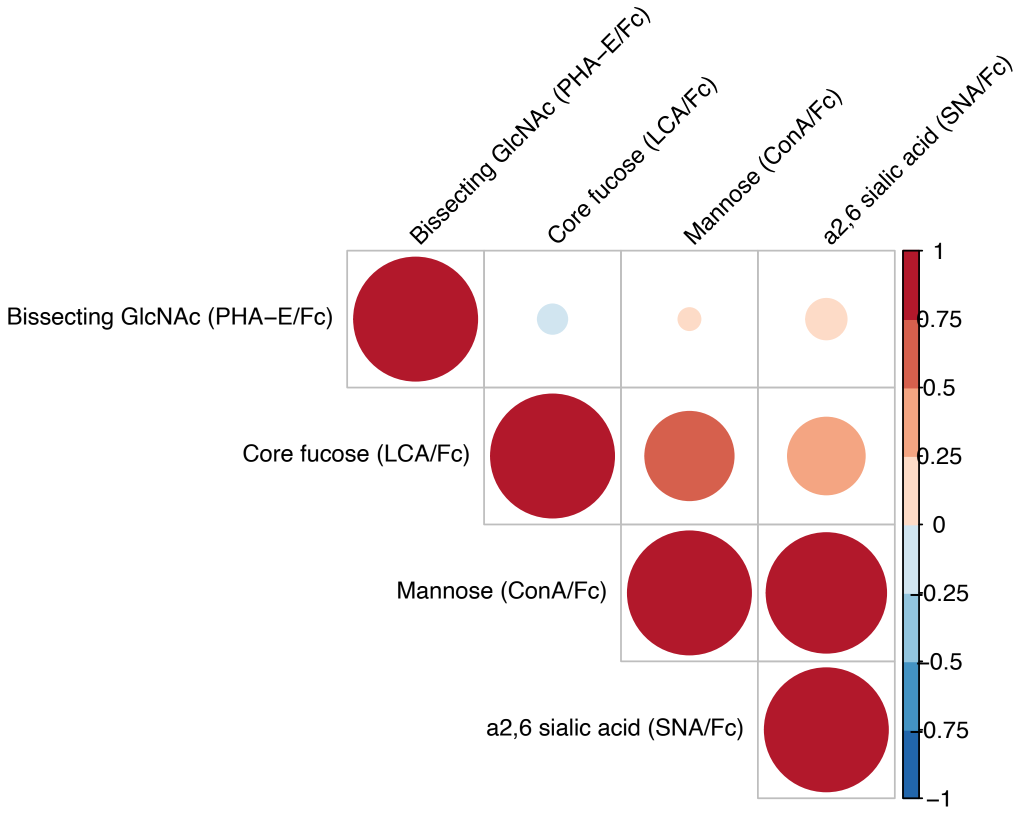


**Supplementary Figure 2: Comparative correlation of Lectin-Binding Profiles Normalized to Fc Levels.** The color of each dot reflects the strength and direction of the correlation (from −1 to 1), with warmer tones indicating stronger positive correlations and cooler tones indicating negative correlations. The size of the dots is proportional to the absolute value of the correlation coefficient, with larger dots representing stronger correlations regardless of direction.

ConA: Concanavalin A; LCA: Lens culinaris agglutinin; SNA: Sambucus nigra agglutinin; PHA-E: Phaseolus vulgaris Erythroagglutinin.

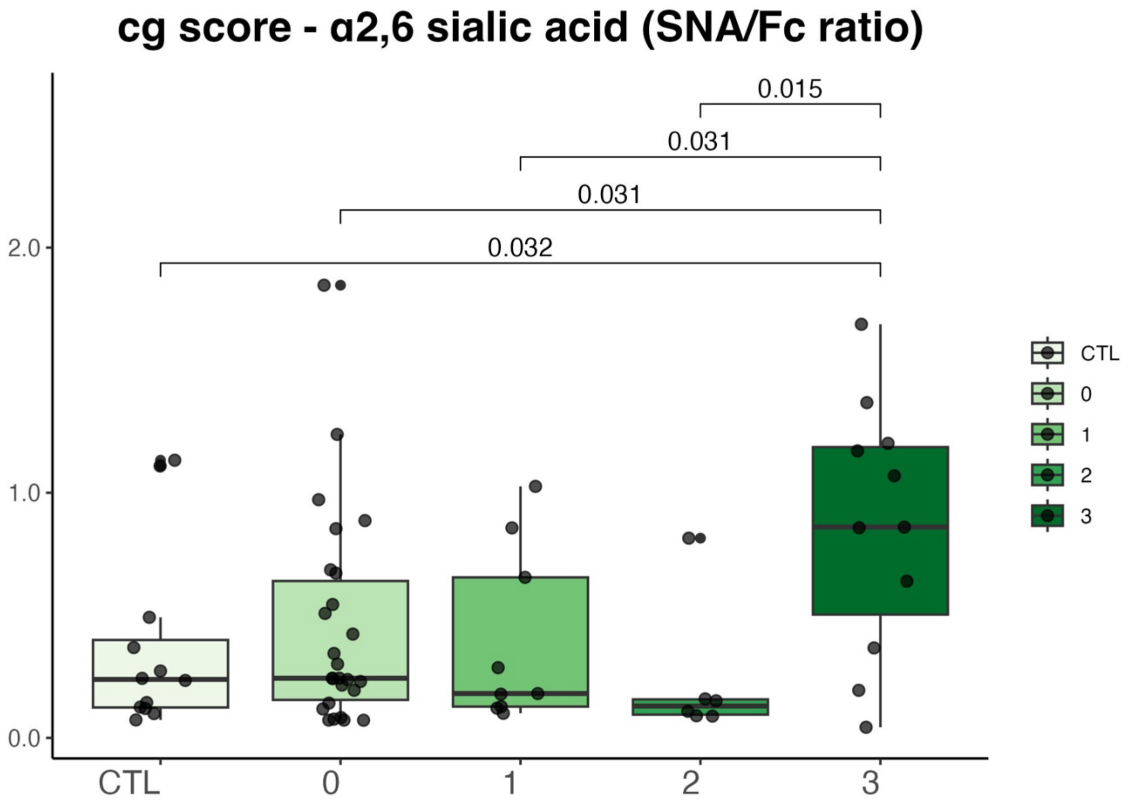


**Supplementary Figure 3: α2,6-Linked Sialic Acid Levels according to CG-Scores.** Boxplot of α2,6-sialic acid (SNA/FC ratio), showing group-specific variations (CTL, and severity of the CG-score).

CTL: control patients, SNA: Sambucus nigra agglutinin.

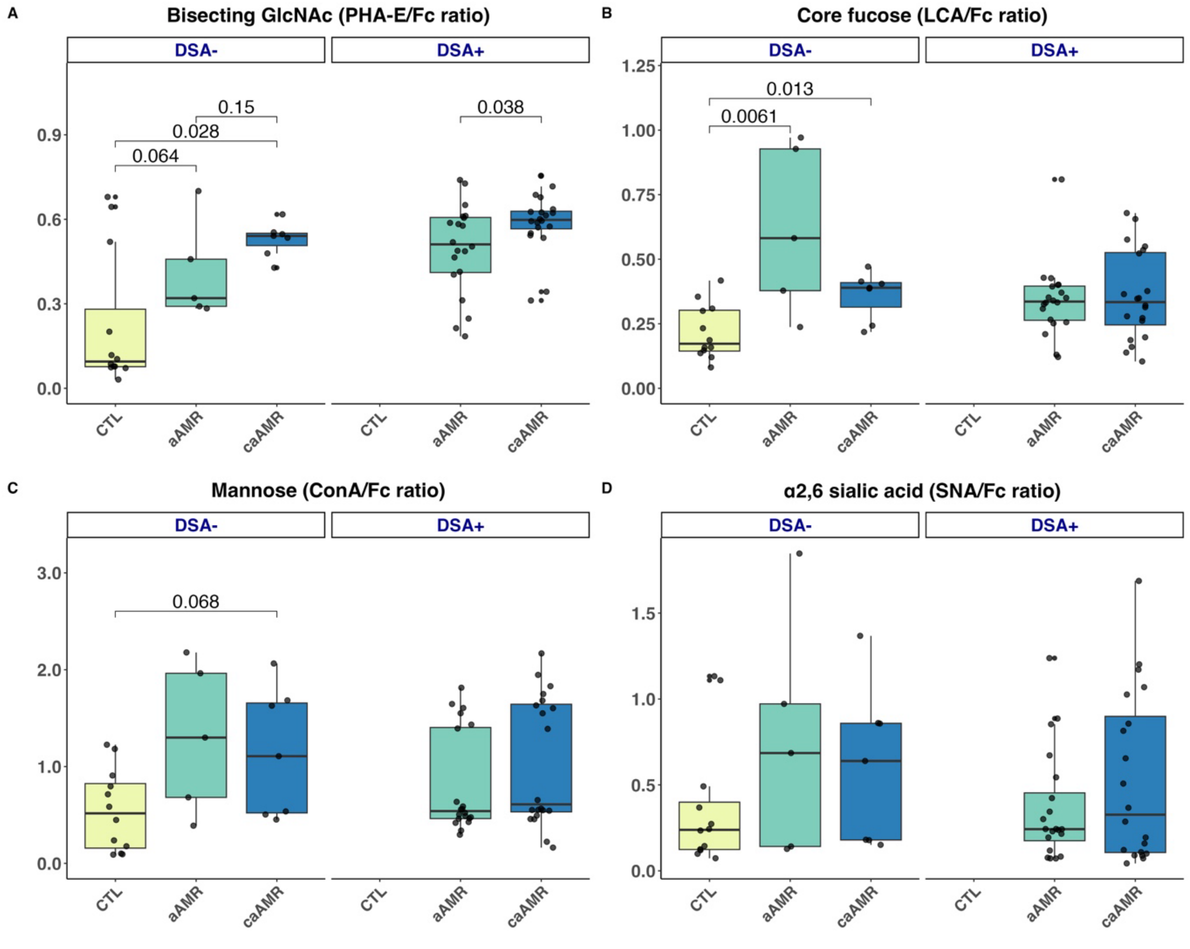


**Supplementary Figure 4: Associations between IgG post-translational modifications and the presence of antibody-mediated rejection according to the presence or absence of Donor-specific antibodies (DSA).** Box plots comparing the amounts of bisecting GlcNAc (PHA-E/Fc ratio in panel A), core fucose (LCA/Fc ratio in panel B), mannose (ConA/Fc ratio in panel C), and α2,6-sialylation (SNA/Fc in panel D) among control patients, patients with acute antibody-mediated rejection and chronic-active antibody mediated rejection, according to the presence or absence of circulating DSA at the time of the biopsy.

aAMR: acute Antibody-Mediated Rejection; caAMR: chronic active Antibody-Mediated Rejection; CTL: control patients, ConA: Concanavalin A; DSA: Donor specific antibodies LCA: Lens culinaris agglutinin; SNA: Sambucus nigra agglutinin; PHA-E: Phaseolus vulgaris Erythroagglutinin.
